# Impaired polyamine metabolism causes behavioral and neuroanatomical defects in a novel mouse model of Snyder-Robinson Syndrome

**DOI:** 10.1101/2023.01.15.524155

**Authors:** Oluwaseun Akinyele, Anushe Munir, Marie A. Johnson, Megan S. Perez, Yuan Gao, Jackson R. Foley, Yijen Wu, Tracy Murray-Stewart, Robert A. Casero, Hulya Bayir, Dwi U. Kemaladewi

**Affiliations:** Div. of Genetic and Genomic Medicine, Dept. of Pediatrics, University of Pittsburgh School of Medicine, Pittsburgh, USA; Dept. of Human Genetics, School of Public Health, University of Pittsburgh, Pittsburgh, USA; Children’s Neuroscience Institute, Dept. of Pediatrics, University of Pittsburgh School of Medicine, Pittsburgh, USA; Sidney Kimmel Comprehensive Cancer Center, Johns Hopkins School of Medicine, Baltimore, Maryland, USA; Dept. of Developmental Biology, University of Pittsburgh School of Medicine, Pittsburgh, USA

**Keywords:** Polyamines, Spermine synthase, spermine, neurological, disease, pathogenesis

## Abstract

Polyamines (putrescine, spermidine, and spermine) are essential molecules for normal cellular functions and are subject to strict metabolic regulation. Mutations in the gene encoding spermine synthase (SMS) lead to accumulation of spermidine in an X-linked recessive disorder known as Snyder-Robinson syndrome (SRS). Presently, no treatments exist for this rare disease that manifests with a spectrum of symptoms including intellectual disability, developmental delay, thin habitus, and low muscle tone. The development of therapeutic interventions for SRS will require a suitable disease-specific animal model that recapitulates many of the abnormalities observed in patients.

Here, we characterize the molecular, behavioral, and neuroanatomical features of a mouse model with a missense mutation in *Sms* gene that results in a glycine-to-serine substitution at position 56 (G56S) of the SMS protein. Mice harboring this mutation exhibit a complete loss of SMS protein and elevated spermidine/spermine ratio in skeletal muscles and the brain. In addition, the G56S mice demonstrate increased anxiety, impaired learning, and decreased explorative behavior in fear conditioning, Morris water maze, and open field tests, respectively. Furthermore, these mice failed to gain weight over time and exhibit abnormalities in brain structure and bone density. Transcriptomic analysis of the cerebral cortex revealed downregulation of genes associated with mitochondrial oxidative phosphorylation and ribosomal protein synthesis. Our findings also revealed impaired mitochondrial bioenergetics in fibroblasts isolated from the G56S mice, indicating a correlation between these processes in the affected mice. Collectively, our findings establish the first in-depth characterization of an SRS preclinical mouse model that identifies cellular processes that could be targeted for future therapeutic development.

## Introduction

The polyamines putrescine, spermidine, and spermine are ubiquitous molecules essential for normal cellular functions. As a group, these polycationic molecules are responsible for maintaining chromatin structure, regulating gene expression, metabolic pathways, and specific ion channels, as well as cell growth and death. Polyamines are also critical contributors to the many processes underlying immune cell activation, wound healing, and general tissue growth and development (1–7). Given their central role in cellular metabolism, the levels of intracellular polyamines are tightly regulated via *de novo* synthesis, interconversion, and transport into and out of the cell (Figure 1).

**Figure 1.**
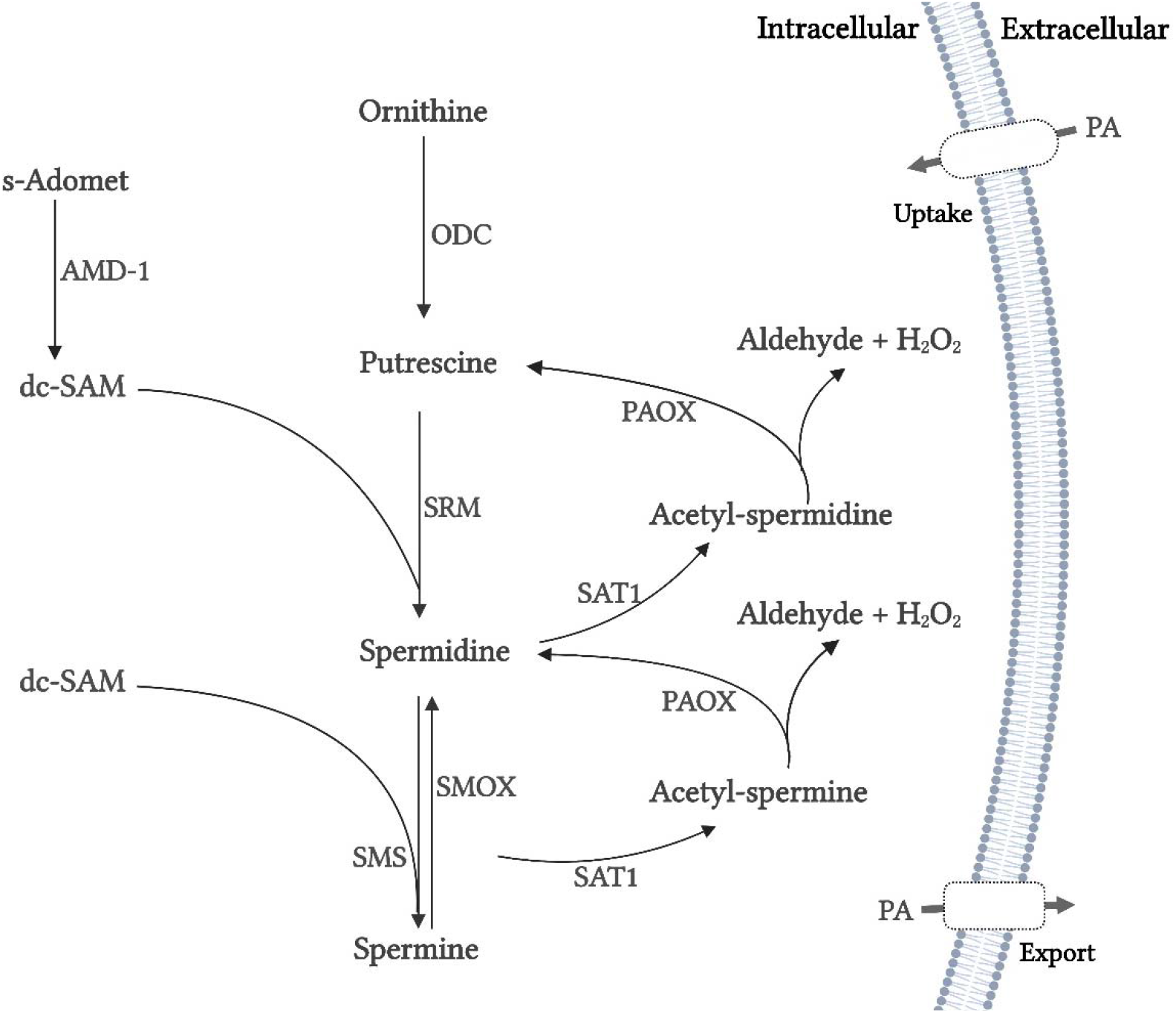
The polyamine metabolic pathway. Abbreviations: ODC, ornithine decarboxylase; SRM, spermidine synthase; SMS, spermine synthase; s-Adomet, s-adenosylmethionine; AMD1, adenosylmethionine decarboxylase; dc-SAM, decarboxylated s-adenosylmethionine; PAOX, acetylpolyamine oxidase; SAT1, spermidine/spermine acetyltransferase; PAs, polyamines.

Most of the intracellular polyamine pool arises from *de novo* synthesis. The diamine putrescine, which is the main precursor in this pathway, is synthesized from ornithine via the actions of the enzyme ornithine decarboxylase (ODC). Spermidine and spermine are higher-order polyamines derived from putrescine. Spermidine synthase (SRM) catalyzes the reaction between putrescine and decarboxylated S-adenosylmethionine (dc-SAM) that generates spermidine. Spermine is then synthesized from spermidine and dc-SAM by spermine synthase (SMS). Spermidine and spermine can be converted back to their respective precursors via the actions of spermidine/spermine N^1^-acetyltransferase (SAT1) and acetylpolyamine oxidase (PAOX). Similarly, spermine oxidase (SMOX) catalysis the direct conversion of spermine to spermidine. Dysregulation or absence of any of these enzymes alters the homeostatic levels of cellular polyamines. Aberrancies in polyamine metabolism have been implicated in numerous diseases, including cancers, Alzheimer’s disease, and Snyder-Robinson Syndrome (SRS) (8–11).

SRS (OMIM: 309583) is a rare (frequency currently unknown, more than 100 reported cases), X-linked intellectual disability syndrome associated with several known deleterious mutations in the *SMS* gene (MIM *300105). These lead to the loss or reduction of spermine synthase enzymatic activity and reductions in the intracellular spermine content of the cell (10). Loss of SMS activity also leads to an accumulation of spermidine and a high intracellular spermidine/spermine ratio. These parameters are use as biomarkers for SRS diagnosis (11). SRS is characterized by intellectual disability, seizures, and general developmental delay (11– 13). Individuals diagnosed with SRS exhibit thin body habitus and low muscle tone (hypotonia). These clinical findings may be apparent at infancy and worsen over time (11,14). Some SRS patients have difficulty walking and may never become ambulatory (12).

Although, a specific mutation in the *SMS* gene was first identified and reported as the cause of SRS as early as 2003 (15), there are no suitable mammalian models to study disease pathophysiology and support the development of effective therapeutic interventions. Previou reports featured a *Drosophila* model in which an appropriate mutation was introduced into th *sms* gene (16) (17). While *Drosophila* models are readily available, many pathogenic factors that are critical to our understanding of vertebrate diseases are not conserved in this species and thus key aspects and mechanisms may not be detectable (17). To address this concern, several groups used the *Gy* (*Gyro*) mice to explore the pathogenesis of SRS (18,19). The *Gy* mouse strain features a complete loss of the chromosomal region containing the *Sms* gene and exhibits several of the abnormalities described in SRS patients, including altered polyamine content, cognitive impairment, and bone abnormalities (20,21). However, in addition to the *Sms* deletion, the *Gy* mice also have a deletion of the *Phex* gene. *Phex* encodes the phosphate-regulating endopeptidase homolog, which is a protein involved in phosphate transport that has been implicated in the pathogenesis of X-linked hypophosphatemia (20,21). These additional *Phex* mutation complicates the interpretation of many of the abnormalities observed in the *Gy* mice, meaning this strain is a poor model to study SRS pathophysiology and for therapy development.

Here, we characterized a recently generated mouse model of SRS (22) that carries a missense mutation in the *Sms* gene (<u>GG</u>C to <u>TC</u>C), a glycine to serine substitution at position 56 of the SMS protein (G56S). This mutation is analogues to that reported in individuals diagnosed with severe SRS (12). In this study, we investigate the disease presentation in the G56S mice and explored potential mechanisms underlying some of the observed abnormalities. Our findings revealed that G56S mice exhibit many of the phenotypic (both behavioral and neuroanatomical) abnormalities reported in SRS individuals (12,23,24). Furthermore, a transcriptomic analysis identified critical changes in gene expression patterns that might contribute to these observed aberrancies.

## Materials and methods

### Mice

All animals used in this study were housed at the University of Pittsburgh Division of Laboratory Animal Resources, Rangos Research Building, following the IACUC protocol number 2206137, which was approved by the University of Pittsburgh’s Institutional Animal Care and Use Committee. The colony of mutant mice was established by breeding female heterozygous *Sms* mutation carriers (C57BL/6J-Sms^em2Lutzy^/J; Jackson Laboratory stock # 031170) and male WT C57BL/6J mice (Jackson Laboratory stock # 000664). The male offspring of this cross that harbored the X-linked G56S *Sms* mutation and WT littermate controls were used in the experiments described in this study. To ensure that only male mice harboring the desired mutation were used, pups were genotyped at Transnetyx.com using the following probes: forward primer ACCTGGCAGGACCATGGATATTTA, reverse primer GTGTTCACATCTAAAGCCCATGAGA, reporter 1 AACAAGAATGGCAGGTAAG and reporter 2 ACGAACAAGAATTCCAGG.

### Open field activity assay

The open field chamber is a hollow square field box equipped with tracking software (ACTITRACK, Panlab/Harvard Apparatus, USA) connected to an infrared tracking system that monitors animal movement. The walls of the box were opacified (covered with aluminum foil) to prevent the environment from influencing the behavior of the mouse undergoing testing. The chamber was divided into two imaginary zones: an outer zone (45 × 45 cm) and an inner or center zone (18.5 cm x 18.5 cm, centered at 22.5 cm from the wall on each side). Experiments were undertaken under constant room temperature (22 – 25ºC) and light levels. The mice were habituated in the procedure room for 15 minutes each time before the assay was initiated. This was done to reduce any stress on the mice before the tests were conducted. Each mouse was released at the same location near the wall of the box and movement was evaluated for 15 minutes using the infrared tracking system. The positions recorded for each mouse were used to generate tracking plots and to determine the distance traveled, speed, and time spent in each zone (i.e., within the entire apparatus and specifically in the center zone). The total amount of time spent and the type of body motion (i.e., rearing, leaning, and vertical activity) detected in the center zone were used as relative measurements of explorative behavior and anxiety-related responses, respectively.

### Auditory-cued fear conditioning

The conditioning procedure was carried out using a specifically designed chamber (model *H10-11M-TC-SF* Coulbourn Instruments, Whitehall, PA, USA). The conditioning chamber (25 × 25 × 25 cm) had three grey methacrylate walls, a grid floor connected to a shock scrambler to deliver foot shock as the unconditioned stimulus (US), and a speaker mounted on the chamber ceiling to deliver audible tones as the conditioned stimulus (CS). The conditioning chamber was fitted with a high-sensitivity camera system that monitored animal movement. The chamber was confined in a ventilated, soundproof enclosure (78 × 53 × 50 cm) on an anti-vibration table in a quiet room. The door to the room remained closed throughout the conditioning and testing periods.

On the first day (fear acquisition), the animals were habituated for 120 sec in the chamber before the delivery of CS-US pairs (i.e., a 75 dB tone [CS] for 20 sec followed by a 15-sec trace and then foot shocks [US] of 0.6 mA for 2 sec) with variable and pseudo-randomly distributed intervals between pairs of stimuli (90 – 203 sec). On the second day (fear retention), the session started with the mice placed in the same environment. During this phase, the mice were provided with no stimulation that might elicit contextual fear responses. Freezing responses in this otherwise familiar environment were monitored.

For the third session, the mice were placed in a different environmental setting (i.e., a chamber with a covered floor and white walls) to assess the retention of cued fear in a novel context. Baseline fear responses were monitored for 90 sec followed by the delivery of three CS (75 dB and 20 s) separated by variable inter-trial intervals (ITIs). The movement of the animal was sampled at a frequency of 50 Hz for quantitative analysis (Freezeframe, Coulbourn Instruments, USA). Freezing was analyzed during the delivery of the CS (20 sec periods) as well as during the 15 sec trace period that would ordinarily precede the US (not delivered) to monitor the associative fear response. The animals were gently handled before, during, and after the test to avoid introducing any additional potential stress before or during each test that could influence the measured responses.

### Morris Water Maze (MWM) Task

The MWM task was performed in a circular pool containing water using the procedure described by Tsien et al. (25) with slight modifications. The animals were trained to find an escape platform that was submerged in the water. The animals were not habituated in the pool before the training. The training protocol (hidden platform, used to evaluate spatial learning) included five sessions with 4 trials per session per day. Navigation was tracked by a video camera and the escape latency (i.e., the time required to locate the platform) was recorded. An animal that failed to locate the platform within 90 sec was guided to the platform. We then performed visible (to measure spatial memory) and probe (to measure non-spatial memory) tests on day six. In the visible test, colored tape was placed at the top of the platform. For the probe test, the platform was removed; the mice were allowed to swim in the pool for 60 □ s, and the time spent in each quadrant of the pool was recorded. The acquired data was analyzed using the ANY-maze software.

### *In vivo* Magnetic Resonance Imaging (MRI) scans

All mice were subjected to *in vivo* brain imaging while under isoflurane anesthesia. The mice were placed in a clear plexiglass anesthesia induction box that permitted unimpeded visual monitoring. Induction was achieved by the administration of 3% isoflurane in oxygen for several minutes. The depth of anesthesia was monitored by toe reflex (extension of limbs, spine positioning) and respiration rate. Once established, the appropriate level of anesthesia was maintained by continuous administration of 1-2% isoflurane in oxygen via a nose cone. The mice were then transferred to the designated animal bed for imaging. Respiration was monitored using a pneumatic sensor placed between the animal bed and the mouse’s abdomen. Rectal temperature was measured with a fiber optic sensor and maintained with a feedback-controlled source of warm air (SA Instruments, Stony Brook, NY, USA).

*In vivo* brain MRI was carried out on a Bruker BioSpec 70/30 USR spectrometer (Bruker BioSpin MRI, Billerica, MA, USA) operating at 7-Tesla field strength and equipped with an actively shielded gradient system and a quadrature radio-frequency volume coil with an inner diameter of 35 mm. Multi-planar T_2_-weighted anatomical images were acquired with a Rapid Imaging with Refocused Echoes (RARE) pulse sequence with the following parameters: field of view (FOV) = 2 cm, matrix = 256 × 256, slice thickness = 1 mm, in-plane resolution = 78 μm X 78 μm, echo time (TE) = 12 msec, RARE factor = 8, effective echo time (ETE) = 48 msec, repetition time (TR) = 1800 msec, and flip angle = 180^0^. Multi-planar diffusion MRI was performed using the following parameters: field of view (FOV) = 2.0 cm, matrix = 128 × 128, slice thickness = 1.5 mm, in-plane resolution = 156 μm X 156 μm, TE = 16.31 msec, TR = 1500 msec, diffusion preparation with the spin echo sequence, diffusion gradient duration = 4 msec, diffusion gradient separation = 8 msec, diffusion direction = 30, number of A_0_ images = 1, and b value = 1500 s/mm^2^.

The MRI data were exported to a DICOM format and analyzed using the open source ITK-SNAP (http://www.itksnap.org) brain segmentation software by 2 independent observers who were blinded to the experimental conditions. The volumes of each region of interest (ROI), including the amygdala, corpus callosum, thalamus, ventricles, hippocampus, and cortex were manually drawn by blinded observers based on the information obtained from the Allen mouse brain atlas (https://mouse.brain-map.org/static/atlas). To account for potential difference in the sizes of brains in G56S and WT mice, volumes from each brain region were normalized to the total brain volume of each mouse.

Diffusion MRI was analyzed by the open source DSI studio (http://dsi-studio.labsolver.org/) to obtain fractional anisotropy (FA). ROIs contributing to quantitative and statistical analyses, including the cortex, hippocampus, thalamus, corpus callosum, and ventricles with cerebrospinal fluid (CSF) were manually segmented and defined by blinded independent observers.

### *In vivo* micro-Computed Tomography (micro-CT) scans

All mice undergoing *in vivo* micro-CT imaging were maintained under general inhalation anesthesia with isoflurane as described for MRI scans above. Once established, anesthesia was maintained with 1.5% isoflurane in oxygen administered using a nose cone and the mouse was transferred to the designated animal bed for imaging. Respiration was monitored as described above. Respiration gating was performed using BioVet system that was triggered by maximal inhalation with a 500 ms trigger delay.

Respiration-gated *in vivo* micro-CT imaging was performed with Siemens Inveon Multimodality micro-CT-SPECT-PET system with the following parameters: full rotation, 360^0^ projections; settle time 1000 msec; 4×4 binning; effective pixel size of 76.75 μm; trans axial field of view (FOV) 78.6 mm with 4096 pixels; axial FOV 76.1 mm with 3968 pixels 80 kV of voltage; current of 500 μA; exposure time of 410 ms. The three-dimensional (3D) micro-CT images were reconstructed using the Feldkamp algorithm and were calibrated in Hounsfield Units (HU). Double distilled water was set at a readout of 0 and air at 1000 HU.

The 3D micro-CT image stacks were analyzed using the Inveon Research Workplace (IRW). The ROI analysis function was used with a thresholding tool that created several ROIs with different Hounsfield Unit (HU). A cylindrical 3D ROI was drawn around the body that encompassed the entire body. All external air around the mouse was excluded from the ROI and a custom threshold was set between 400 – 5700 HU to capture the bones. The mean HU values obtained from each ROI were used to quantify bone density.

### Body composition measurements

Body composition (percentage lean and fat weight) of the mice was measured by quantitative MRI (EchoMRI, Echo Medical Systems, Houston, TX). Animals were placed in thin-walled plastic cylinders with plastic restraining inserts. Each animal was briefly subjected to a low-intensity electromagnetic field that measured total body composition. Percentages of fat and lean weights were determined based on total body weight.

### Primary cell isolation

Primary fibroblast cells from the ears of the G56S and WT mice were isolated using the protocol described by Khan and Gasser (26).

### RNA isolation and quantitative polymerase chain reaction (qPCR)

Total RNA was isolated from mouse tissues using the Nucleospin RNA Plus kit (Macherey-Nagel, cat# 740984.50), following the manufacturer’s instructions. cDNA synthesis was performed using the iScript Reverse Transcriptase Supermix kit (BioRad, Cat# 1708841) according to the manufacturer’s instructions. qPCR was performed using 2X SYBR Green Fast qPCR Mix kit (ABclonal, cat# RM21203) in a C1000 Touch Thermal Cycler (BioRad, USA). The primer sequences used to amplify target genes of interest are listed in Supplementary **Table 1**. The expression of endogenous *Gapdh* was used as an internal control to measure the relative expression of genes of interest. The 2^ΔΔ*C*t^ was used to assess relative fold change in gene expression in tissue samples from WT and G56S mice. Values are presented as the percentage change in fold expression.

### RNA-sequencing (RNA-seq) and pathway enrichment analysis

After completing the RNA extraction procedure described above, samples were submitted to the Health Sciences Genomic Core at the UPMC Children’s Hospital of Pittsburgh. RNA quality was determined using the Agilent Bioanalyzer 2100 (Agilent Technologies, USA). cDNA libraries were prepared using a 3′-Tag-RNA-Seq library kit (Illumina). Sequencing was performed using one lane of a Hi-Seq 4000 platform with pair-end 40 bp reads. Analysis of sequence reads, including quality control, mapping, and generation of tables of differentially expressed genes (DEGs), heatmaps, and volcano plots) were performed using the Qiagen licensed CLC Genomic Workbench software version 22.0.1. Pathway enrichment analysis of the DEGs was performed using the Qiagen-licensed Ingenuity Pathway Analysis (IPA) software. The gene expression profile identified by RNA-seq was validated by qPCR as described above.

### *In vivo* analysis of mitochondria respiration

Oxygen consumption rates (OCRs) were determined with a Seahorse XFe96 Extracellular Flux Bioanalyzer (Agilent Technologies, Santa Clara, California, USA). Primary fibroblasts plated in 96-well assay plate at a density of 40,000 cells/well were cultured overnight and then equilibrated with Seahorse XF base medium (Agilent Technologies) supplemented with glucose, sodium pyruvate, and L-glutamine at 37°C in a CO_2_- free incubator for 1 hour prior to the assay. Mitochondrial function was assessed by sequential addition of 1.5 μM oligomycin, 1 μM FCCP (carbonyl cyanide-4-[trifluoromethoxy] phenylhydrazone), and 0.5 μM rotenone/antimycin A by the Seahorse Bioanalyzer. Data was normalized by the total protein content of the cells.

### Protein isolation, quantification, and western blotting

Total proteins were extracted from tissues isolated from G56S and WT mice tissues using RIPA homogenizing buffer (150 μL of 50 mM Tris HCl pH 7.4, 150 nM NaCl, 1 mM EDTA) followed by homogenization using a bullet blender. After homogenization, 150 μL of RIPA double-detergent buffer (2% deoxycholate, 2% NP-40, 2% Triton X-100 in RIPA homogenizing buffer) supplemented with protease inhibitor cocktail (Roche, cat# A32953) was added to the tissue homogenate followed by incubation on a shaker for 1 h at 4ºC. The tissue homogenate was then centrifuged at 11,000 *g* for 10 min at 4ºC. The resulting supernatant was used to quantify total protein using the Pierce BCA protein assay kit (Thermo Scientific, cat# 23225) according to the manufacturer’s protocol. Twenty micrograms of total protein were fractionated on 4 – 12% gradient gel (Thermo Scientific, cat# NP0336BOX). After proteins had separated on the gel, they were transferred by electroblotting onto a polyvinylidene fluoride (PVDF) membrane and blocked with 5% non-fat milk in TBS-Tween-20. The membrane was then incubated overnight with rabbit anti-spermine synthase (Abcam, cat# ab156879 [EPR9252B]) or rabbit anti-vinculin (Abcam, cat# ab129002 [EPR8185]). After incubation with the primary antibody, the membranes were washed and then incubated with the secondary antibody (Goat Anti-Rabbit IgG – HRP conjugate, Bio-Rad cat# 1706515) for one hour at room temperature. Specific protein bands were detected using SuperSignal™ West Femto Maximum Sensitivity Substrate (Thermo Scientific, cat# 34095). Bands corresponding to immunoreactive SMS and Vinculin were identified and quantified using the ChemiDoc Imaging System (BioRad).

### Polyamine measurement

The polyamine content in isolated tissues was measured by the precolumn dansylation, high-performance liquid chromatography method described by Kabra et al. using 1,7-diaminoheptane as the internal standard (27).

### Statistical analysis

Statistical analysis was performed using Graphpad Prism software vs 9.0. Each variable was statistically compared between the WT and G56S mice using unpaired Student’s t-test unless otherwise stated. A *p* value less than 0.05 was considered statistically significant.

## RESULTS

### *Sms* mutation altered polyamine content in mouse tissues

To determine whether the substitution of two nucleotides (<u>GG</u>C to <u>TC</u>C) in exon two of the *Sms* gene altered its expression profile and tissue polyamine levels, we measured the mRNA and protein levels of SMS in tissues isolated from both the G56S and WT mice. While there were no significant changes in the level of *Sms* mRNA (Figure 2a), we observed a near-complete loss of SMS protein in both the brain and skeletal muscles of G56S mice (Figure 2b). Similarly, HPLC analysis of tissue polyamines revealed elevated levels of both putrescine and spermidine and a significant decrease in spermine in brain tissue from G56S mice. The spermidine to spermine ratio was 7-times higher in brain tissue from G56S mice compared to WT controls (Figure 2c). Our findings also revealed a significant increase in spermidine but decrease spermine content in skeletal muscles of the G56S strain (Figure 2d and inset). Putrescine was below the limits of detection in the sampled muscle. Figure 2e presents the two-dimensional (2D) crystal structure of the dimerized and fully functional SMS protein including the C-terminal (catalytic) and N-terminal (dimerization) domains; the amino acid within the latter domain at position 56 (WT, glycine) is highlighted in Figure 2f. Figure 2g presents the mutant SMS protein with a serine at position 56; the extended side chain characteristic of this amino acid may interfere with monomer dimerization, as previously described by Zhang et. al., (28) and may lead to the loss of SMS protein despite normal transcript level.

**Figure 2.**
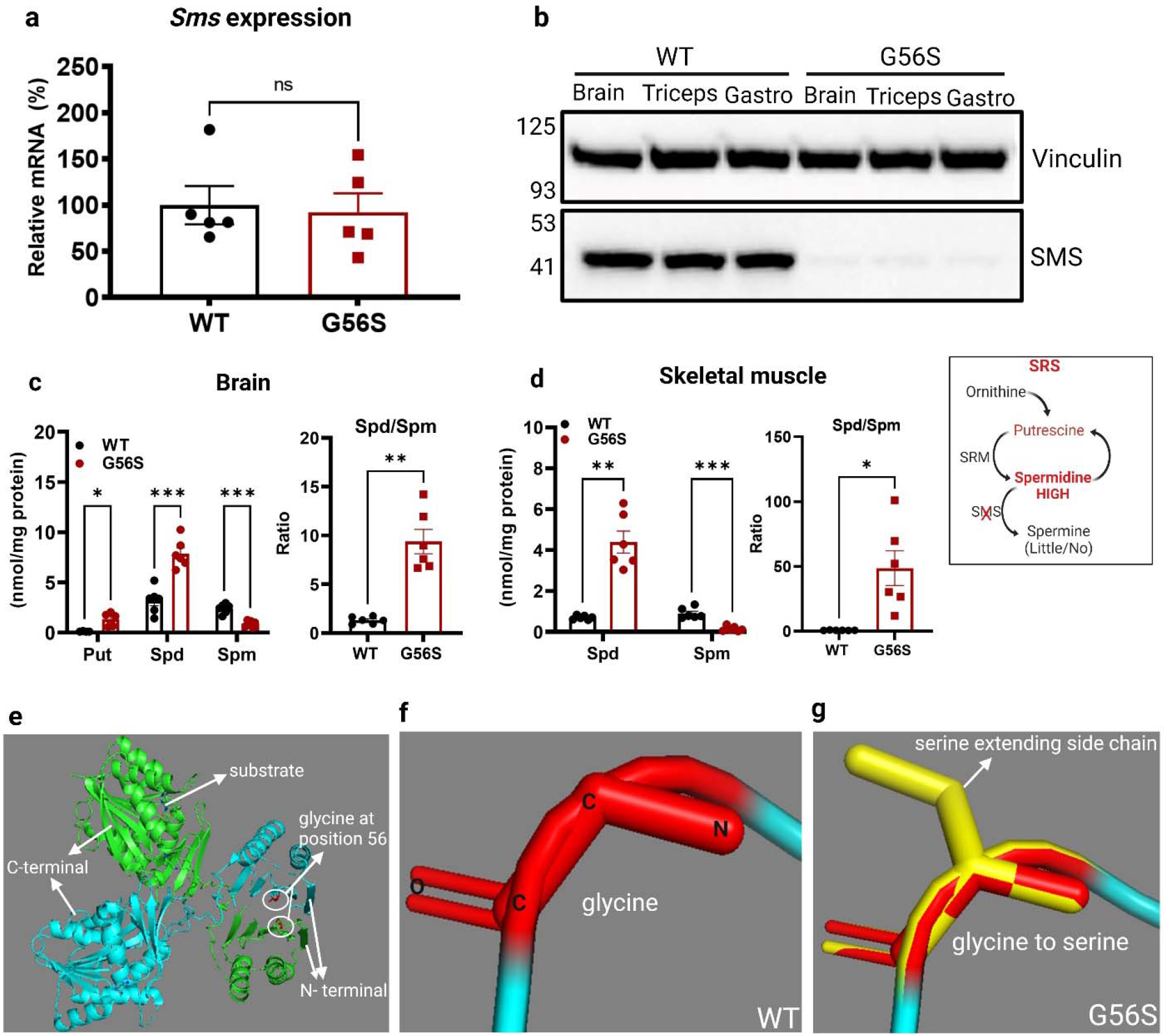
Tissue polyamine content and a potential mechanism underlying SMS protein loss. (a) Mouse brain *Sms* gene transcript level as quantified by RT-qPCR. The level of gene transcript in the mutant was compared to the WT mice set at 100% reference value (b) Expression of SMS protein in isolated brain and skeletal muscles (triceps and gastrocnemius) from WT and G56S mice. (c) Determination of brain polyamine content and SPD/SPM ratios in WT and G56S mice quantified by HPLC. (d) Skeletal muscle polyamine content in both G56S and WT mice. Note: spermine levels were below the limit of detection in the G56S skeletal muscle. (e) 2D-crystal crystal structure of the SMS protein with glycine at position 56 in the N-terminal region (circled). The 2D-crystal structure of SMS was modeled from protein data bank ID: 3C6M. (f) The atomic structure of glycine at position 56 in the N-terminal region of SMS protein. (g) Serine in place of glycine at position 56 of SMS protein; the extended serine sidechain is highlighted in yellow. Panels a-d, values shown are mean ± standard error of the mean (S.E.M.); n = 3 – 5 mice per group; \**p < 0*.*05*, \*\**p < 0*.*01*, ns = not significant.

### Biometric parameters are significantly altered in G56S mice

Patients diagnosed with SRS typically exhibit an asthenic physique with thin body build and short stature. Many of these patients also exhibit significant bone abnormalities and low muscle tone (11). To understand the mechanisms contributing to these physical aberrations, we determined the impact of altered polyamine content on the growth and development of G56S mice. Routine measurements revealed that the G56S mice have significantly lower body weight than age-matched WT counterparts and also gained little to no weight throughout the duration of the study period (Figure 3a). Similarly, the G56S mice exhibit significantly reduced body lengths (Figure 3b). Collectively, these results suggest that the mutant mice may experience failure to thrive compared to their WT littermates. Also, an analysis of the body composition of mutant mice revealed a higher percentage of lean weight and a lower fat weight compared to their WT counterparts (Figure 3c). This finding may reflect the diminished body build typically observed in patients diagnosed with SRS. Similarly, quantification of bone density by micro-CT revealed that the G56S mice exhibited decreased bone density compared to their WT littermates (Figure 3d). Taken together, this data suggests that changes in polyamine content observed in the G56S mice tissues have an impact on overall development and result in general failure to thrive.

**Figure 3.**
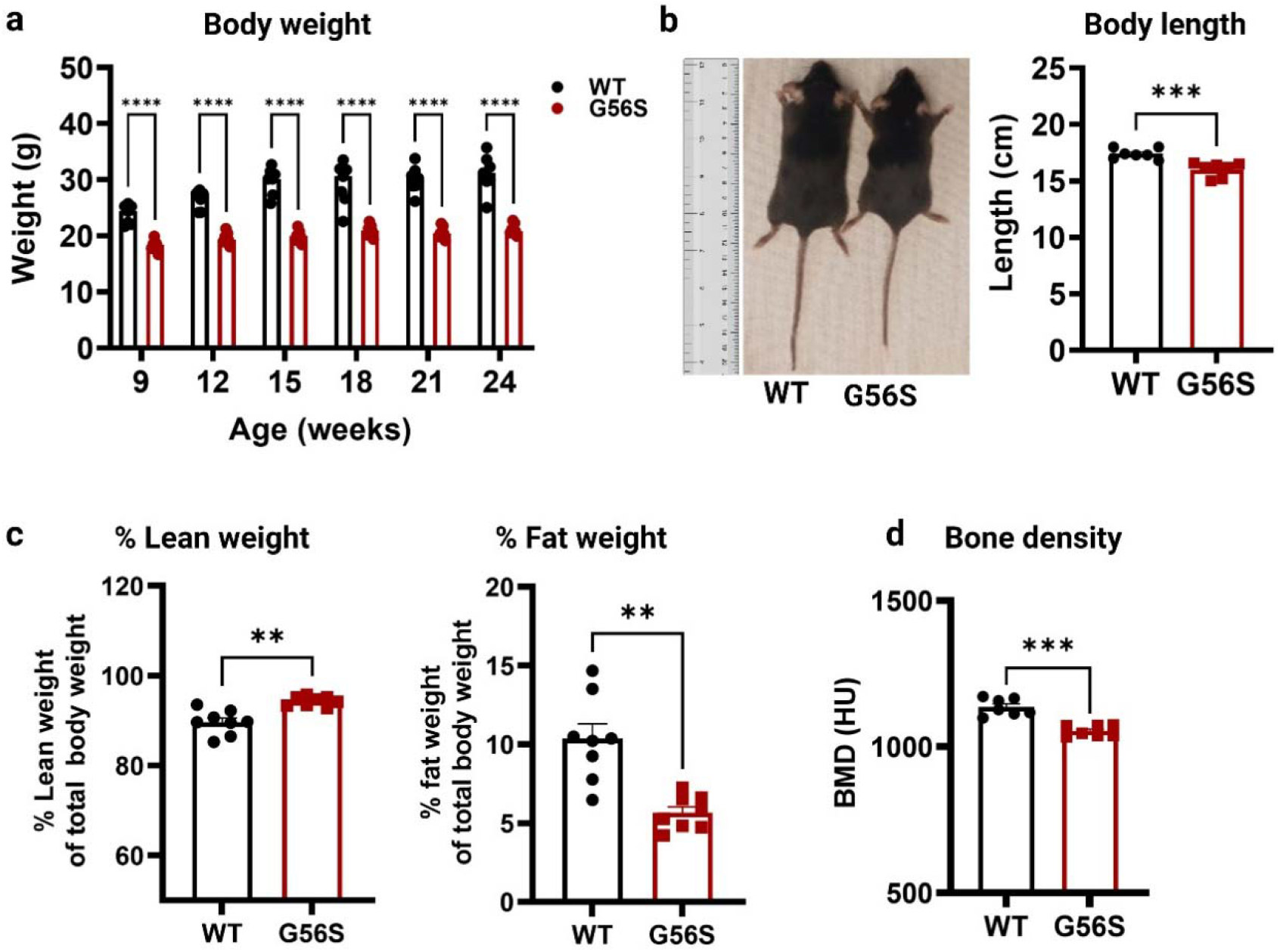
Biometric analysis of male G56S and WT mice. (a) Total body weight was measured in 3-week intervals. (b) Body length of 24-week-old mice. (c) Body composition of 15-week-old mice (% lean and % fat weight) determined by by Echo-MRI scan. (d) Bone mineral density (BMD) of 20-week-old mice measured by micro-CT scan. Values shown are mean ± S.E.M., n = 7 mice per group, ** *p* < *0*.*01*, *** *p < 0*.*001*, **** *p < 0*.*0001*.

### G56S mice exhibit signs of cognitive impairment

Mild to severe cognitive impairment is one of the major clinical observations of SRS. Two patients diagnosed with the G56S mutation were reported to have severe cognitive disabilities (12). To explore the impact of the G56S mutation on cognition, we evaluated the performance of G56S and WT mice using the Morris water maze (MWM) test. For this test, the mice were placed in a pool of water and were subjected to five days of training in which they learned to locate an escape platform that was submerged and thus hidden under the water (Figure 4a, panel 1). The time taken to find the platform on each day was recorded.

**Figure 4.**
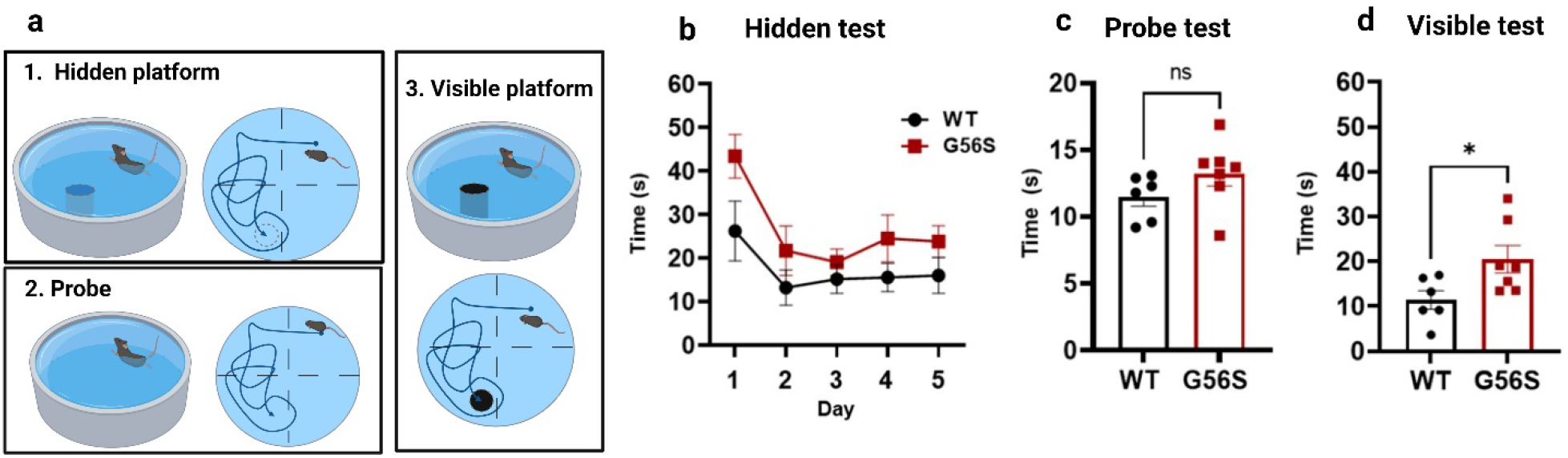
Performance of G56S and WT mice in a Morris Water Maze (MWM) test. (a) Description of the three components of the MWM test. (b) Time required to locate a hidden escape platform on each day of a five-day training period. (c) Time spent in the escape quadrant (probe test without the platform) given a limit of 60 sec on day 6. (d) Time required to locate a visible escape platform on day 6. Data are mean ± S.E.M., n=7 mice per group; two-way ANOVA for repeated measures (a) and unpaired *t*-tests for (b) and (c), not significant (ns); \**p < 0*.*05*.

While our findings revealed no significant differences in the time required to find the escape platform, we observed a trend that suggests that the WT were somewhat more effective than the G56S mice at performing the task (Figure 4b). A secondary probe test, in which the platform was completely removed from the pool (day 6; see Figure 4a, panel 2) revealed no significant differences in the time spent in the escape quadrant (Figure 4c), however, the G56S mice required significantly more time to locate a visible platform (see Figure 4a, panel 3) compared to the WT mice (Figure 4d). This result may indicate that the WT mice learned and retained the information needed to complete this task more effectively than the G56S mice. Overall, the results of the MWM test suggest that the G56S mice exhibit relatively mild learning impairments compared to the WT mice.

### Diminished explorative behavior in an open field test was observed in the G56S mice

Anxiety-related responses are among the major symptoms of many neurological and neurodevelopmental disorders, including SRS. To study this behavior, an open field tests can be employed to evaluate and characterize different mouse strains. In our study, both the WT and G56S mice were provided the opportunity to explore an open field arena which had been divided into an outer and inner zones (Figure 5a). We assessed the total activity, the resting time in each zone, the number of times each mouse entered the inner zone and the time spent in the inner zone. The test was performed once every three weeks for 15 min to characterize changes in behavior with advancing age.

**Figure 5.**
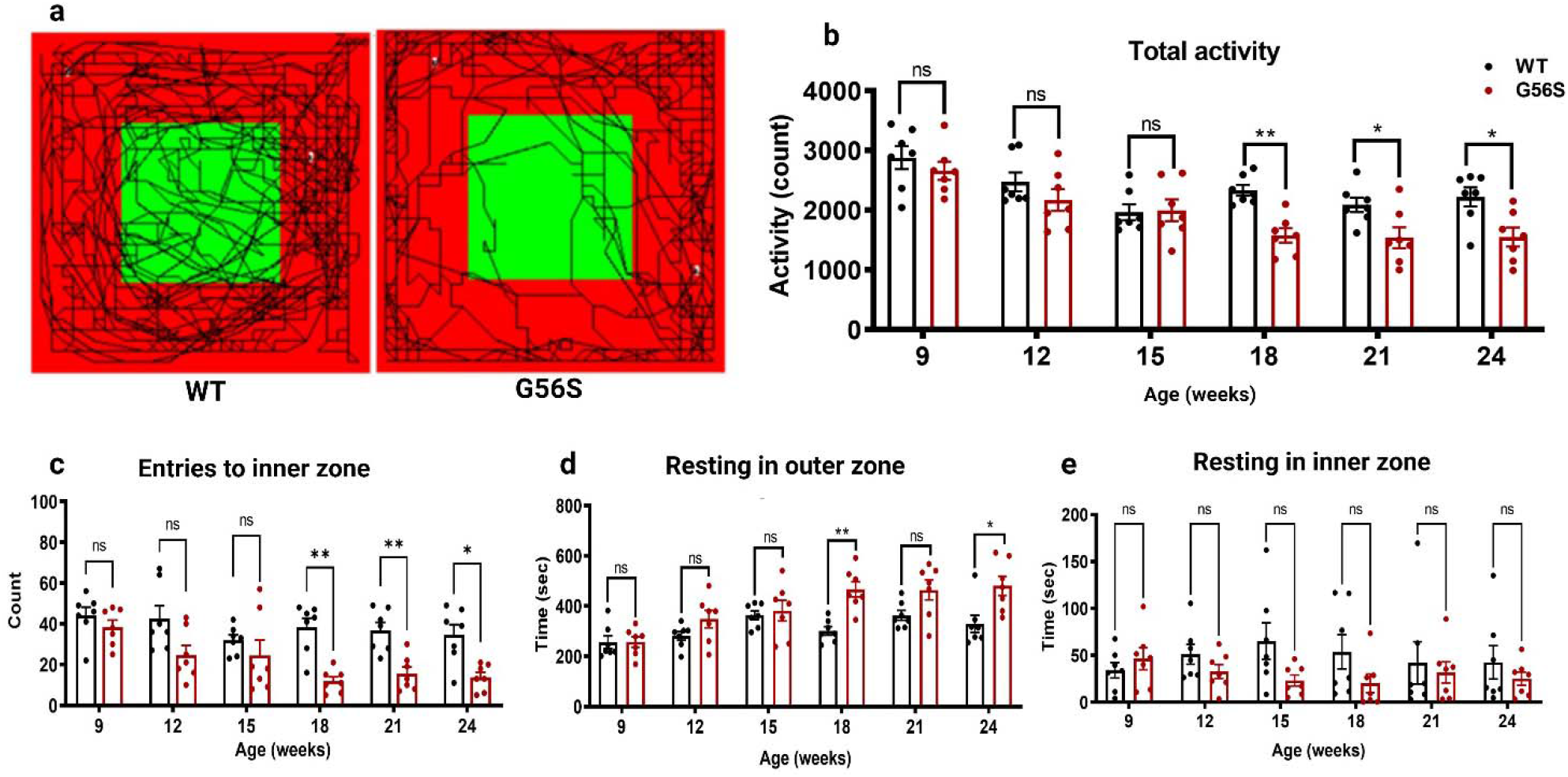
Anxiety-related response monitoring in an open field test. (a) Movement pattern of the animals for 15 min in the open field test at 24 weeks of the age. (b) Total activity of the animals in the open field test measured as the number of total numbers of movement, rearing, and other bodily activities. (c) Number of entries to the center zone of the open field test. (d and e) Resting time of the animals in the outer and center zones of the open field respectively. Data shown are mean ± S.E.M., n=7, \**p* < 0.05; \*\**p* < 0.01; ns, not significant.

Among our results, we found that the G56S mice were less active compared to their WT counterparts (Figure 5a). While no statistically significant differences were detected between the two strains when the mice were less than 18 weeks old, the G56S mice became significantly less active with increasing age compared to their WT littermates (Figure 5b). Similarly, G56S mice were much less likely to enter the inner zone of the open field arena than their WT counterparts beginning at 18 weeks of age (Figure 5c). These results suggest the possibility that the G56S mice experience slow but steady disease progression.

Older G56S mice also spent significantly more time resting in the outer zone (Figure 5d) and less time resting in the inner zone of the open field compared to the WT control (Figure 5e). Collectively, these findings suggest that G56S mice exhibit less explorative behavior than their WT counterparts and that these responses may represent higher anxiety or fear that increases as the mice age.

### Fear-related responses were higher in the G56S mice

Fear-related responses exhibited by both WT and G56S mice were assessed via measurements of stress-induced freezing. This response is an innate anti-predator fear-related behavior that is characterized by a complete tonic immobilization while sparing respiration. This test uses auditory-cued fear conditioning, which requires the mouse to associate an aversive outcome with an otherwise unrelated cued stimulation.

In this experiment, anxiety or fear responses are expressed as the percentage of time spent in a freezing position after sound stimulation (an auditory cue) followed by foot shock (aversive condition).

On the first day of the test, the mice were trained to associate the sound (conditioned stimulus, CS) with the foot shock (unconditioned stimulus, US). The percentage of time spent in stress-induced freezing was recorded during the CS and after delivery of the US (Figure 6a). We observed a consistent increase in the freezing responses following the CS in both groups of mice; this response reached a plateau after the delivery of the fourth sound stimulation and decreased after the fifth (Figure 6b). These results suggest similar rates of fear acquisition by both groups of mice. However, we observed significant differences in the freezing responses displayed during the inter-trial intervals (ITIs, i.e., the period between the last foot shock and the next sound stimulation). The G56S mice exhibited significantly longer, and more frequent freezing responses compared to the WT mice (Figure 6c). Similarly, WT mice recovered ambulation more rapidly than the G56S mice following administration of the CS and US. This result indicates that the G56S mice exhibit more profound fear responses following stimulations than their WT counterparts.

**Figure 6.**
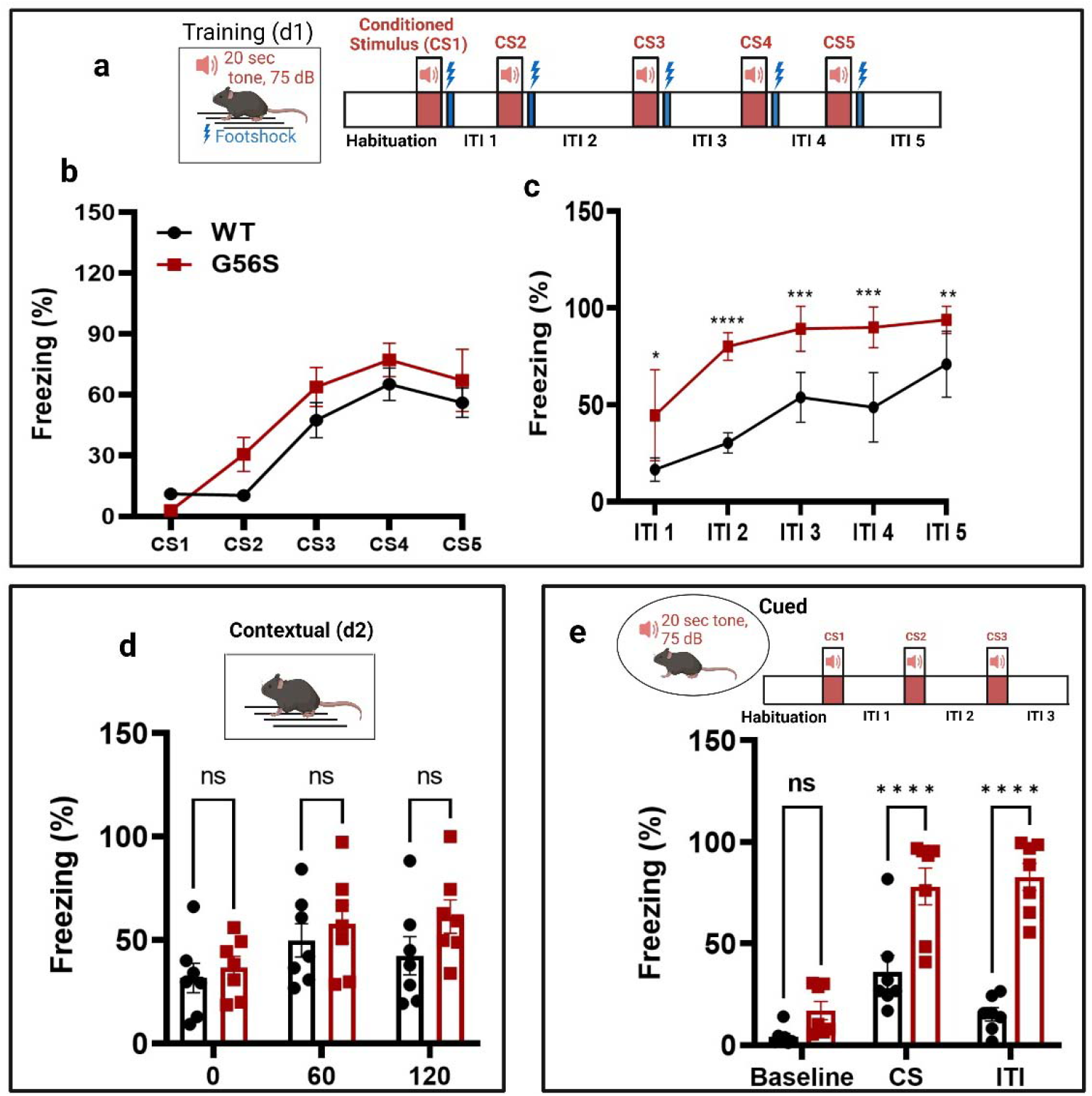
Auditory-cued fear responses following a conditioned stimulus (CS). (a and b) Fear learning and fear response levels after each conditioned stimulus (CS; inter-trial interval, ITI) were assessed in 4–5-month-old G56S and WT mice (n=7 per group). The fear response was expressed as the percentage of time spent in a stereotypical freezing state during the presentation of the CS repeated five times during fear acquisition (CS 1-5; a 75 dB tone lasting 20 sec followed by a foot shock). This was repeated four times with staggered inter-trial intervals. The inset (a) documents the experimental sequences on day 1, which include habituation (hab), tone presentation (CS, loud-speaker symbol), shock delivery (lightning symbol), and ITIs of varying durations. (c) WT and G56S mice demonstrated significantly different responses during fear acquisition (genotype effects and genotype trial interactions were evaluated by two-way ANOVA for repeated measures). (d) Fear response in contextual setting, without sound and shock stimulation, performed 24 hours after fear acquisition. (e) Cue-fear response (with a changed environment) induced by CS alone (tone: 20 sec, repeated three times with no foot shock) with variable ITIs. The Baseline value represents the percentage spent in the freezing state during habituation before the CS (sound); the CS response represents the average of three CS trials; the bars labeled ITI represent the average of the three it is for each mouse. The inset documents the experimental sequences used on day 2, including habituation (hab), tone presentation (CS, loud-speaker symbol), and ITIs of varying durations. Results obtained from WT and G56S mice were compared using unpaired *t*-tests. Values shown are mean ± S.E.M., \**p* < 0.05; ** *p* < 0.01; \*\*\**p* < 0.001; \*\*\*\**p* < 0.0001.

We then compared the freezing responses of G56S and WT mice in a contextual test. This test was performed 24 hours after the CS-US training and involved no stimulation; the mice were placed in the same experimental chamber and their freezing responses in this environment were measured. While no statistically significant differences were observed, we detected a pattern that suggested that the G56S might exhibit increased freezing responses compared to the WT mice (Figure 6d). This trend suggests that the innate fear responses exhibited by the G56S mice may be more profound when compared to those of their WT littermates.

In the cued test, we evaluated the freezing responses of both WT and G56S mice to the CS only in a different environmental setting. After measuring baseline freezing responses during an initial 60 sec habituation period, the mice were subjected to 20 sec of auditory stimulation (CS). Their fear response during the CS and various intertrial intervals were then monitored. While the G56S mice exhibited comparatively higher baseline freezing responses compared to their WT counterparts in the new environmental setting, the differences did not achieve statistical significance. By contrast, the G56S mice exhibited significantly higher fear responses compared to the WT controls both during the CS as well as the ITIs (Figure 6e). Of note, the percentage freezing during the ITIs exhibited by the G56S mice was not only elevated, but it also remained at a level similar to that observed during the CS. Taken together, these data suggest that the G56S mice exhibit higher anxiety-related fear responses than their WT counterparts. This finding may represent a specific neurological dysfunction similar to those observed in patients diagnosed with SRS.

### G56S mice exhibit neuroanatomical changes

We explored brain anatomical structures to determine whether the G56S mutation and resulting alterations in polyamine metabolism were associated with major structural changes. We also determined how altered brain structures might correlate with the behavioral defects described in previous tests. For these experiments, brain volumes were assessed using T2-weighted anatomical scans and diffusion tensor imaging (DTI) (Figure 7a). Other DTI parameters collected included fractional anisotropy (fa), mean diffusivity (md), axial diffusivity (ad), and radial diffusivity (rd). The results of whole-brain imaging showed that the G56S mice have significantly smaller brain volumes than WT mice (Figure 7b). Similarly, analysis of several specific brain regions that were selected based on reports describing SRS patients (23,24) and our behavioral data revealed that the volumes of the amygdala (involved in fear learning and emotional responses), the hippocampus (involved in cognitive functions), and the corpus callosum were all significantly lower in the G56S mouse strain (Figure 7c). Also, DTI analysis of the various brain regions revealed that fa, which is a measure of the microstructural integrity of the white matter of the amygdala and the corpus callosum, was significantly lower in the G56S mice compared to the WT controls. The other regions, including the hippocampus and the cortex, exhibited decreasing trends, although they were not statistically significance (Figure 7d).

**Figure 7.**
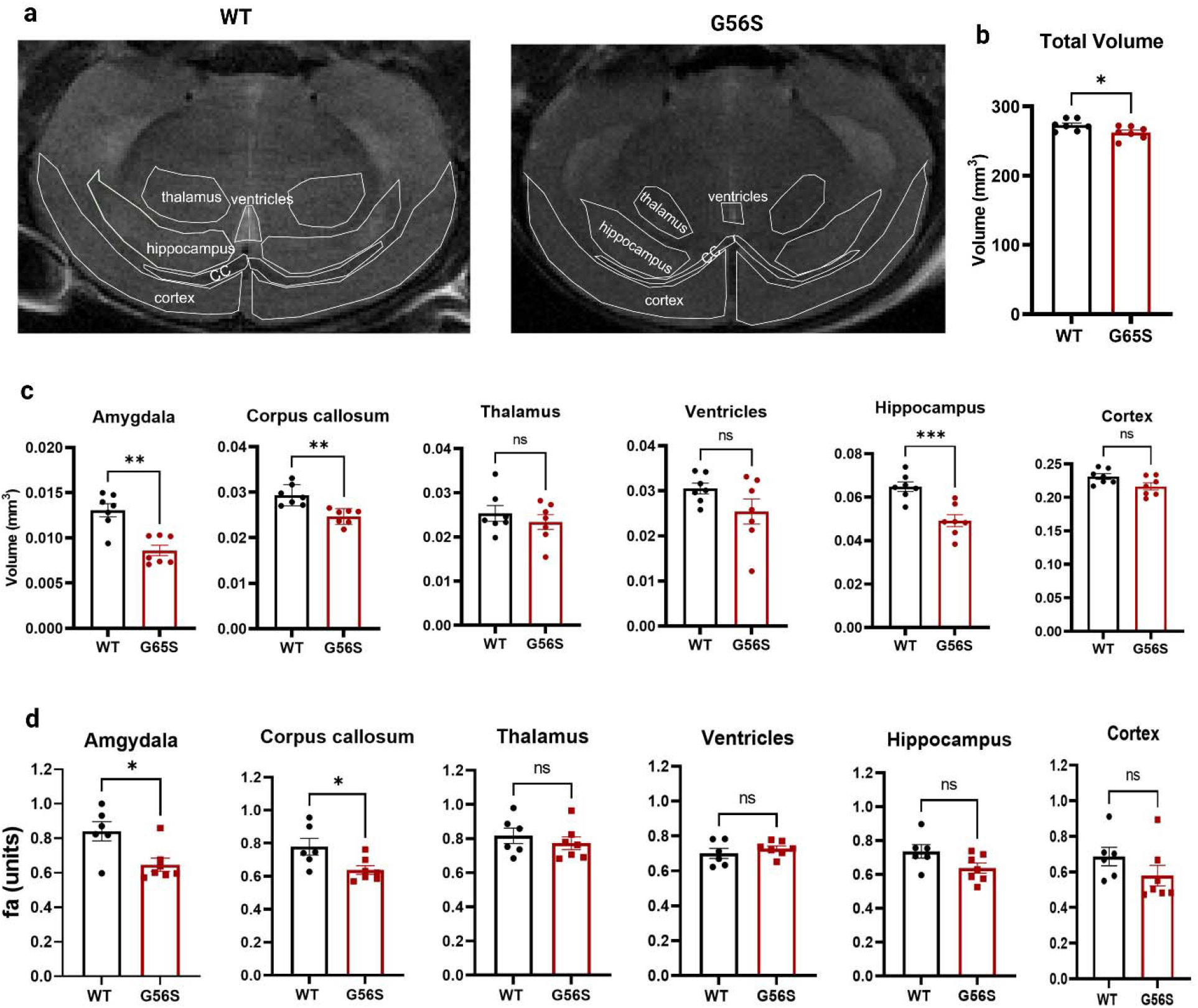
*In vivo* MRI structural analyses. (a) Representative MRI images of coronal sections of brains from WT and G56S mice. Annotations of different regions of the brain were based on the Allen Mouse Brain Atlas. (b and c) MRI volumetric analyses of the total brain volume and volumes of annotated regions highlighted in panel (a). Quantification of the volume of each region was normalized to the total brain volume for each sample. Volumes of each brain region were quantified using ITK snap software to assess RER8 MRI scan images. (d) Fractional anisotropy (fa) was quantified using DSI studio software. Comparisons of single variables between WT and G56S mice were performed using unpaired *t*-tests. Data shown are mean ± S.E.M., n= 5-7, \**p* < 0.05; \*\**p* < 0.01; \*\*\**p* < 0.001; ns, not significant. **Note**: The amygdala region was not visible in the image shown in panel (a) above.

### Mutation in *Sms* gene alters transcriptomic profile of the G56S mouse brain cortex

To elucidate the molecular mechanisms underlying some of the observed phenotypic and behavioral differences, we isolated RNA and performed a transcriptomic analysis of the brain cortex tissue of WT and G56S mice. We focused on the cortex because of its role in directing higher complex tasks, including learning, memory, and consciousness. Furthermore, results from previous studies suggest that spermine may have a protective role specifically within the cerebral cortex (23,29). The results of our transcriptomic analysis of brain cortex tissue from WT and G56S mice revealed more than 1,000 differentially expressed genes (DEGs) (Figure 8a and Supplementary **Table 2**). A heatmap revealed differential expression of genes involved in several key cellular and metabolic processes (Figure 8b).

**Figure 8.**
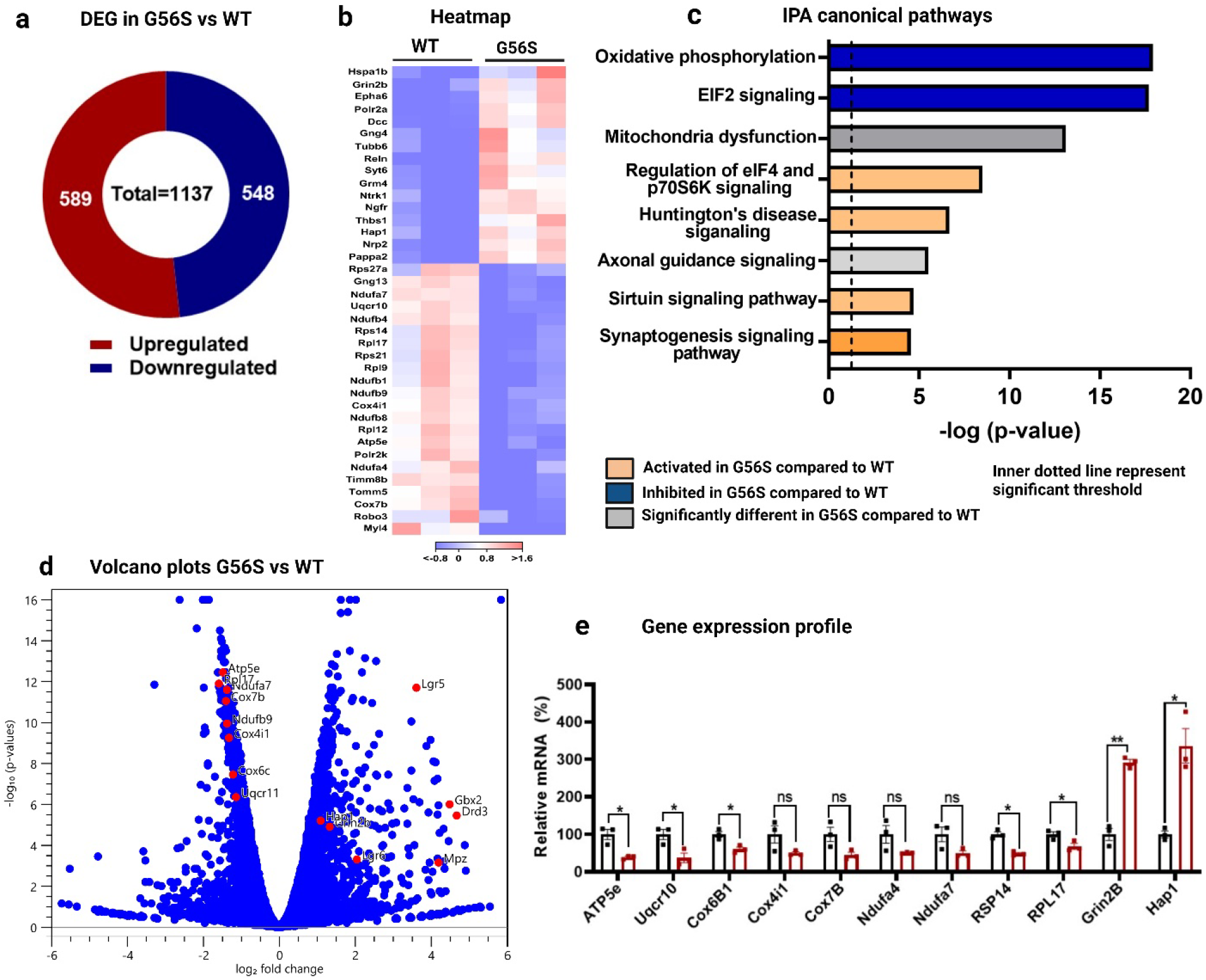
Transcriptomic analysis of isolated brain cortical tissues. (a) Differentially expressed genes (DEGs) were identified using Qiagen CLC Genomic Workshop software. (b) Heatmap of selected genes that exhibit statistically significant differences in expression (*p < 0*.*05*) and absolute values of log_2_-fold change (LFC) greater than or equal to 1. (c) Gene enrichment analysis of up- and downregulated transcripts with *p < 0*.*05* and absolute LFC ≥ 0.5 in the brain tissue from G56S compared to WT mice. Gene enrichment analysis was done using the Qiagen Ingenuity Pathway Analysis (IPA) software. (d) A volcano plot showing the relative expression of selected genes involved in oxidative phosphorylation and other pathways based on the results of the enrichment analysis. (e) qPCR validation of RNA-seq results for selected genes implicated in oxidative phosphorylation, eukaryotic initiation factor 2 (eIF2) signaling, and Huntington’s disease. The data shown mean ± S.E.M, (n=3 for both WT and G56S mice). \**p* < 0.05; \*\**p* < 0.01, ns, not significant.

Importantly we found differential expression of genes that contribute to mitochondrial function and ribosomal protein synthesis. We performed gene enrichment pathway analysis to identify possible metabolic pathways that might be altered in the G56S mice. Our results revealed downregulation of pathways involved in mitochondrial oxidative phosphorylation (OXPHOS) and ribosome protein synthesis (i.e., eukaryotic initiation factor 2 [eIF2] signaling) in the G56S mice (Figure 8c). In addition, Huntington’s disease, sirtuin, and synaptogenesis signaling pathways were all upregulated in the G56S mice brain cortex compared to the WT (Figure 8c). The expression pattern of genes involved in these metabolic processes were further confirmed in the G56S mice relative to the WT by a Volcano plot (Figure 8d). We observed decreased expression of *ATP5e, Uqcr10, Cox6B1, Cox4i1, Cox7b, Ndufa4, and Ndufa7* (all involved in mitochondria OXPHOS), as well as decreased expression of *Rpl17* and *Rsp14* (both implicated in ribosome protein synthesis via eIF2 signaling) in the brain cortex of the G56S mice compared to the WT control (Figure 8e). Furthermore, we found increase expression of *Hap1* and *Grin2b*, which are Huntington-associated protein 1 and ionotropic NMDA receptor subunit 2b respectively (Figure 8e), both implicated in Huntington disease signaling.

Collectively, these data strongly suggest that the G56S mutation in the *Sms* gene and the subsequent alterations in brain polyamine content result in changes in the expression profile of genes involved in central cellular and metabolic processes of the brain. These observations may explain one or more of the phenotypic abnormalities observed in G56S mice.

### G56S mice exhibit impaired mitochondrial respiration

To confirm the downregulation of mitochondrial oxidative phosphorylation predicted by gene expression analysis, we isolated primary fibroblasts from both the WT and G56S mice and evaluated mitochondrial respiration using the XFe96 Seahorse bioanalyzer. Our results revealed that both basal and oligomycin-sensitive respiration were significantly diminished in fibroblasts isolated from G56S mice (Figures 9a and 9b). Similarly, maximum respiration and rates of ATP synthesis were also significantly reduced in fibroblasts from the G56S mice compared to the WT (Figures 9c and 9d). Taken together, these data suggest that SMS deficiency and impaired polyamine metabolism in G56S fibroblasts will lead to impaired mitochondrial bioenergetics and functions.

**Figure 9.**
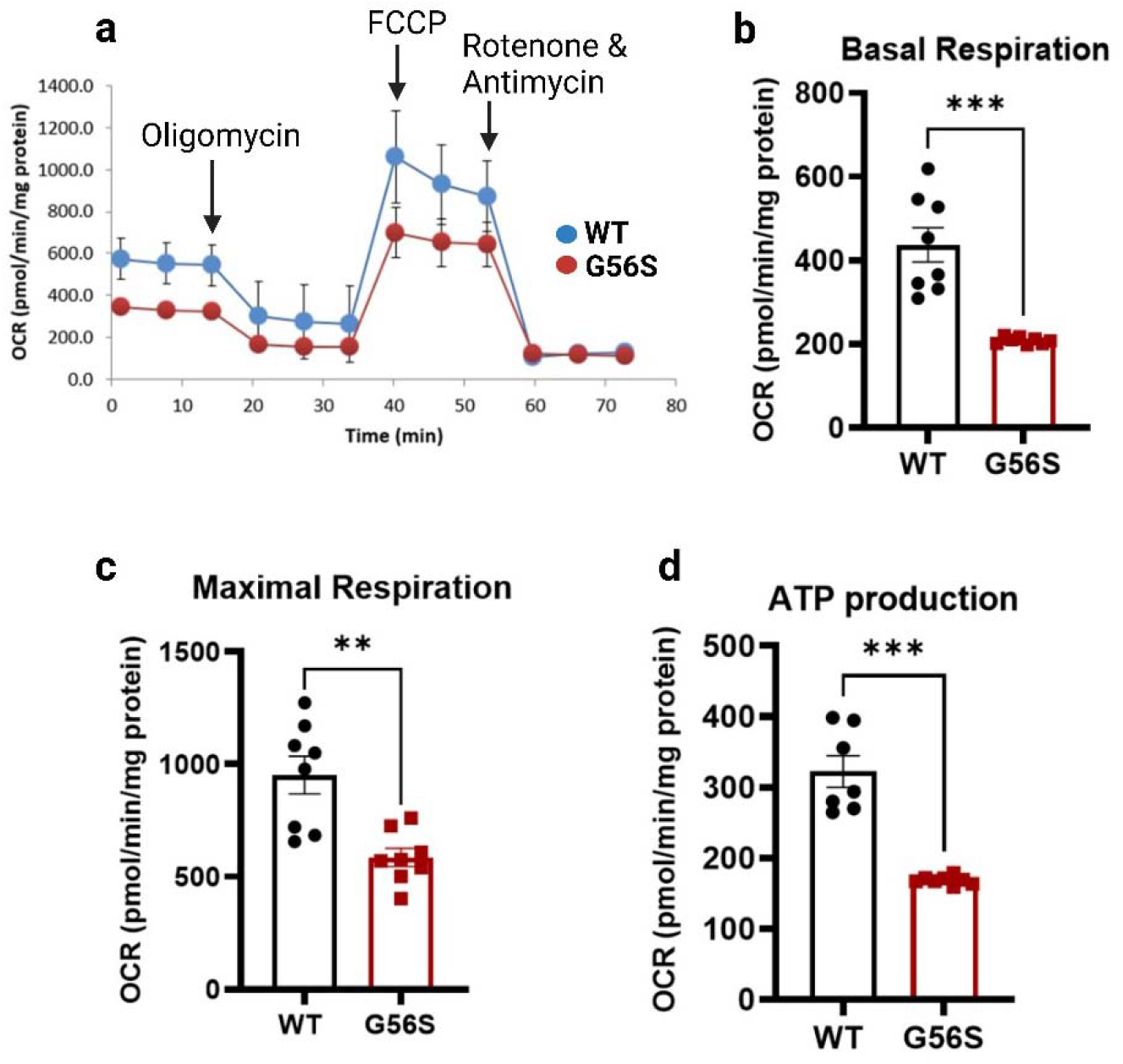
Mitochondrial respiration in isolated fibroblasts. (a) Respiratory profiles of primary fibroblasts isolated from WT (blue line) and G56S (red). Oligomycin (ATP synthase inhibitor), FCCP (H^+^ ionophore), and rotenone/antimycin (mitochondria complex I/III inhibitors) were added at the times indicated. (b) Basal respiration, (c) maximal respiration, and (d) ATP production in fibroblasts from WT and G56S mice were assessed using a Seahorse XFe96 analyzer. Data shown are mean ± s.e.m. of average n=8 technical replicates of two independent experiments. *\*\*p < 0*.*01; ***p < 0*.*001*.

## Discussion

One of the main challenges of preclinical research intended to elucidate the mechanisms underlying diseases associated with cognitive impairment is the identification of a robust and translationally relevant animal model. While the G56S mouse features an *Sms* gene-specific missense mutation that replicates a genetic lesion identified in patients with severe forms of SRS, it is not known whether this mouse represents a useful tool for therapy development. Therefore, the goal of this study is to provide a detailed behavioral and neuroanatomical assessment of these mice as a baseline for future phenotype reversal studies. In addition, we studied the transcriptomic landscape of the brain cortex to identify gene expression patterns that may contribute to the abnormalities observed in the mice and possibly in SRS patients.

SRS is a disease characterized by abnormal somatic features and a general failure to thrive. Patients are described as having asthenic body builds, mild short stature, and diminished body mass. These patients also exhibit abnormal bone structure and sustain frequent fractures (11). In the current study, we report that the G56S mice exhibit lower body weights as well as reduced overall length, bone mineral density, and body fat composition compared to their WT counterparts. These findings suggest that the G56S mouse model reproduces many of the abnormal clinical features described for SRS. However, the impact of the resulting disruptions in polyamine metabolism and their contributions to disease-specific symptomatology remain unknown. A previous study that focusing on bone marrow-derived multipotent stromal cells (MSCs), revealed that the mRNA silencing of *SMS* gene resulted in impaired cell proliferation and a reduced capacity for osteogenesis (30). These results suggest a possible link between the *Sms* mutation, dysregulated polyamine metabolism, and the decreased bone mineral density observed in both patients and the G56S mice. While most SRS patients, including those with the G56S mutation (12) are eventually diagnosed with kyphoscoliosis, no scoliosis was detected in micro-CT scans of these mice. However, we cannot rule out the possibility that abnormal spines may develop in older mice.

The decreased body fat composition might be attributed to the increase in tissue spermidine, as this polyamine has been implicated in promoting lipolysis (31). However, it is also possible that the absence of SMS can result in impaired mitochondrial functions (16,30). In this case, the mice will depend more heavily on glycolysis as a means of energy generation; resulting in the utilization of greater amounts of food with less available to be converted to and stored as body fat. The decreased body weight seen in these mice supports the general idea that disturbances in polyamine homeostasis impair cell growth and tissue development (4) which may lead to general growth failure.

Similar to what has been reported in many SRS patients, the G56S mice display signs of cognitive impairment and exhibit significant reduction in exploratory behavior when evaluated in an open field test. These findings suggest that these mice exhibit amplified anxiety-related behaviors. This hypothesis was further studied using the fear-conditioning test. Mutant mice displayed significantly heightened fear responses when presented with an auditory conditioned stimulus (CS) followed by unconditioned stimulus (US; foot shock). The increase in fear responses in both the contextual and cued tests further confirm the observed heightened anxiety-related behavior in the G56S mouse strain. Overall, this data provides some evidence suggesting the existence of neurological abnormalities in the mutant mice. Since polyamines are important contributors to the development of the nervous system (32), several regions of the brain might be contributing to these behavioral defects. Both the amygdala and the hippocampus in the G56S mice, which are brain regions involved in fear-associated memory and learning showed decrease volume compared to the WT, similar to that reported in SRS patients (23). Thus, these results suggest that impaired polyamine metabolism and excess spermidine accumulation may lead to atrophy and neuronal loss in these regions (23) as well as the defects in behavioral and learning outcomes observed in these mice. The possibility that disrupted polyamine metabolism might lead to brain atrophy was further confirmed by the decrease in the fa value found in G56S mice. Although we did not measure the polyamine content in specific regions of the brain, we believe that the total brain polyamine content most likely reflects the overall content in the different regions, given that G56S mice lack SMS in all tissues and that this mutation is not tissue or brain-region specific.

Another potential mechanism that might explain the behavioral defects observed in G56S mice is spermidine-mediated disruption of receptor signaling. In an earlier study, Rubin *et al*. (33) reported that intra-amygdala administration of spermidine in an experimental rat model resulted in a dose-dependent increase in freezing responses. These results suggested that the excess accumulation of spermidine in the brains of G56S mice might contribute to the observed increase in anxiety-related behaviors. The mechanisms underlying spermidine-mediated increases in fear responses have not yet been clarified.

Spermidine may regulate the function of the amygdala via interactions with and modulation of the ion channel receptor for N-methyl-D-aspartate (NMDA); an earlier report detailed polyamine-mediated negative regulation of this receptor (34). Administration of arcaine, a putative competitive antagonist at the polyamine binding site of the NMDA receptor, resulted in a decrease in spermidine-induced fear responses in rats (33). Collectively, these results suggest that spermidine levels may have an impact on amygdala function and that excess accumulation of spermidine may induce a fear response as well as other behavioral abnormalities seen in the G56S mice. It is important to note that anxiety has been identified as one of the main symptoms of numerous neurological disorders (35). Although there are no clinical reports that document this specific behavior in SRS patients, family members have confirmed anxiety and the prevalence of fear-related behaviors in some patients (personal communication, Snyder-Robinson Foundation Conference 2022).

In addition to the neuroanatomic defects, other potential mechanisms contributing to the phenotypic abnormalities observed in the G56S mice were revealed by the transcriptomic analysis. These include impaired mitochondrial function, alterations in ribosomal protein synthesis signaling pathways, and upregulation of genes implicated in the pathogenesis of Huntington’s disease. The mitochondria are important energy-generating cellular organelles; mitochondrial dysfunction has been implicated in a variety of different neurological or neurodegenerative diseases (36). Results from previous studies have suggested that impaired mitochondrial function might contribute to the pathogenesis of SRS (16,30) via one of several potential mechanisms:

1. Increased spermidine levels that accumulate in cells that lack SMS may promote the synthesis and release of reactive oxygen species (ROS) secondary to increased catabolism. Elevated ROS results in mitochondrial oxidative stress and impaired mitochondrial function (16).
2. Polyamines play essential roles in modulating gene expression. Earlier reports suggest that spermine modulates mammalian mitochondrial translation initiation processes (37,38). Thus, the lack of SMS or spermine could inhibit synthesis of mitochondrial proteins and result in impaired mitochondrial functions.
3. Normal mitochondrial metabolism can result in the accumulation of potentially damaging levels of by-products including ROS and Ca^2+^ (39). As a polycationic molecule, spermine can scavenge mitochondrial ROS (40,41), and reduce the levels of mitochondrial permeability transition pore (mPTP) generated in response to Ca^2+^ accumulation (42). Thus, the lack of SMS or an observed decrease in cellular spermine content may result in mitochondrial damage.
4. Another potential mechanism of mitochondrial impairment in SRS may relate to the decreased expression of nuclear genes encoding mitochondrial proteins reported in this study. Although we do not yet understand how *Sms* mutations and/or decrease spermine content result in the changes in gene expression pattern observed in this study, either factor may be involved in direct or indirect interactions with critical transcription factors. It will be important to identify relevant transcription factors as this may improve our understanding of how spermine and/or SMS modulate mitochondrial function.

In conclusion, efforts to develop effective therapies for SRS will require a better understanding of the disease pathophysiology as well as suitable disease-specific animal models that recapitulate many of the critical abnormalities diagnosed in these patients. The findings presented in this study suggest that the G56S mouse is a good model that can be used to study SRS pathogenesis and may be an important tool for therapeutic development. This study also provides parameters that may be used to assess the effectiveness of therapy for SRS in a murine model. Finally, the data shown here offers insight that could be used to improve the current clinical management of SRS.

## Acknowledgements

We thank the Snyder-Robinson Foundation, C. Lutz, and A. Zuberi for generating and providing the G56S mouse model; A. Schmidt, E. Goetzman, and K. Schwab for the advice and assistance with animal imaging; W. MacDonald, R. Elbakri, and A. Chattopadhyay for the help on transcriptomic and pathway analyses; C. van ‘t Land, A. Karunanidhi, and G. Vockley for SeaHorse experiment; O. Tomisin for help with the 2D protein crystal structure modeling. Members of the Kemaladewi laboratory are acknowledged for technical support and input in this study. This research was supported in part by the University of Pittsburgh Center for Research Computing through the resources provided. Specifically, this work used the HTC cluster, which is supported by NIH award number S10OD028483. This work was supported by the RK Mellon Institute for Pediatric Research Postdoctoral Fellowship (to O.A.), the Chan Zuckerberg Rare-as-One Initiative/Snyder-Robinson Foundation and University of Pennsylvania Orphan Disease Center Million Dollar Bike Ride (MDBR-20-135-SRS) to R.A.C and T.M.S., and the Children’s Trust of UPMC Children’s Hospital of Pittsburgh Foundation, Dept. of Pediatrics, Univ. of Pittsburgh School of Medicine, Research Advisory Committee Grant, NIH Director’s New Innovator Award DP2-AR081047, NIH R01-AR078872 (to D.U.K).

## Author contributions

Conceptualization: O.A. and D.U.K; Investigation: O.A., A.M., M.J., M.S.P., Y.G., Y.W., J.F., T.M.S, R.A.C.; Formal analysis and visualization: O.A., A.M., M.E.P., Y.G., Y.W., H.B., D.U.K; Writing: O.A. and D.U.K.; Supervision, D.U.K. All authors read and commented on the manuscript.

